# From Fossils to Living Canids: Two Contrasting Perspectives on Biogeographic Diversification

**DOI:** 10.1101/2023.08.30.555443

**Authors:** Lucas M. V. Porto, Renan Maestri, Thijs Janzen, Rampal S. Etienne

**Affiliations:** Ecology Department, Universidade de São Paulo, São Paulo, SP, Brazil; Ecology Department, Universidade Federal do Rio Grande do Sul, Porto Alegre, Rio Grande do Sul, Brazil; Groningen Institute for Evolutionary Life Sciences, University of Groningen, Groningen, The Netherlands

**Keywords:** diversification rates, fossil information, lineage dispersal, species selection, trait inheritance

## Abstract

The Canidae are an ecologically important group of dog-like carnivores that arose in North America and spread across the planet around 10 million years ago. The current distribution patterns of species, coupled with their phylogenetic structure, suggest that Canidae diversification may have occurred at varying rates across different biogeographic areas. However, such extant-only analyses undervalued the group’s rich fossil history because of a limitation in method’s development. Current State-dependent Speciation and Extinction (SSE) models are (i) often parameter-rich which hinders reliable application to relatively small clades such as the Caninae (the only extant subclade of the Canidae consisting of 36 extant species); and (ii) often assume as possible states only the states that extant species present. Here we extend the SSE method SecSSE to apply to phylogenies with extinct species as well (111 Caninae species) and compare the results to those of analyses with the extant-species-only phylogeny. The results on the extant-species tree suggest that distinct diversification patterns are related to geographic areas, but the results on the complete tree do not support this conclusion. Furthermore, our extant-species analysis yielded an unrealistically low estimate of the extinction rate. These contrasting findings suggest that information from extinct species is different from information from extant species. A possible explanation for our results is that extinct species may have characteristics (causing their extinction), which may be different from the characteristics of extant species that caused them to be extant. Hence, we conclude that differences in biogeographic areas probably did not contribute much to the variation in diversification rates in Caninae.

## 1. INTRODUCTION

From a single ancestor around 40 million years ago (Mya), the Canidae family became one of the most widespread and ecologically diverse groups among Carnivora, inhabiting several distinct environments, and being present in all continents, except in Antarctica (Wang and Tedford 2008; Prothero 2013). Canids originated in North America, where successive radiation events gave rise to three subfamilies, Hesperocyoninae, Borophaginae and Caninae (Wang et al. 2004). The first two subfamilies were endemic to North America, and went extinct without reaching other continents, before the geological events that connected N. America to Eurasia and South America around 11 Mya (Geffen et al. 1996; Cox 2000; Macdonald, D.W & Sillero-Zubiri 2004; Wang and Tedford 2008; Potter and Szatmari 2009), i.e., the uplifts of the Bering Strait and the Isthmus of Panama (MacNeil 1965; Hopkins 1967; Montes et al. 2015). Caninae was the only subfamily that managed to cross the land bridges and disperse from North America, producing more than a hundred species, of which 36 are still extant today (Porto et al. 2019).

Although Caninae have a rich fossil history, raw fossil locations paint a limited picture of biogeographic events. Most of what we know is related to the geographic location from which lineages originated (both fossil and extant species), and the most likely routes that canids used to disperse (Wang and Tedford 2008; Porto et al. 2021). Our understanding on how biogeographic events shaped the evolutionary dynamics of canids is therefore still limited. More precisely, we still lack a complete understanding of how dispersal events to new continents affected Caninae diversification rates. Pires et al. (2015) demonstrated how continental dispersals affected the evolution of carnivores in general, but focused only on dispersal between Eurasia and North America, disregarding the rest of the planet. If invasions of new areas have an impact on canid diversification rates, we might expect a scenario of ecological opportunity (EO) upon the arrival in a new continent lacking competitors (Simpson 1953).

Several studies have indicated that dispersal events to new areas can trigger exceptional shifts in species diversification by EO (Mahler and Losos 2010; Mahler et al. 2010; Yoder et al. 2010; Algar and Mahler 2016). Geographic colonization of new areas will lead to range expansion and establishment of new populations, which is likely to increase the rate of speciation (Cardillo et al. 2005; Rabosky and Glor 2010; Etienne et al. 2012; Uribe-Convers and Tank 2015; Kennedy et al. 2017). Local extinction, however, leads to range contraction implying that the greater the range, the smaller the chance of complete extinction of the species (McKinney 1997). It is evident that species diversification can be linked to distinct factors, however it is not always clear how speciation, extinction and dispersal can be disentangled. Over the last few years, several studies have attempted to fully integrate phylogenetic comparative methods together with ecologically relevant traits to understand how biodiversity can be generated (Mairal et al. 2015; Pires et al. 2015, 2017; Herrera-Alsina et al. 2018; O’Donovan et al. 2018). A promising way to such integration is through diversification methods that estimate rates of speciation, extinction and dispersal.

The state-dependent speciation and extinction (SSE) family of models (Maddison 2006; FitzJohn 2010, 2012) was developed to elucidate the impact that trait changes have on patterns of lineage diversification. In an explicit geographical scenario, the GeoSSE (geographic state-dependent speciation and extinction) model (Goldberg et al. 2011) would be ideal to test the influence of geographic distribution on Caninae diversification, but it was recently found that the initial SSE models have a high Type I error in detecting the influence of traits on diversification rates, as they attribute rate variation directly to trait influences without allowing for rate variation being due to hidden factors (i.e. a null model) (Fitzjohn 2012; Machac 2014; Rabosky and Goldberg 2015). To avoid false positives, Beaulieu and O’Meara (2016) proposed the HiSSE model (hidden-state-dependent speciation and extinction), which can be used to detect if diversification events are more related to an unknown hidden trait than to the observed character. GeoHiSSE (Caetano et al. 2018), a combination between GeoSSE and HiSSE models, focused on geographical state dependence in particular. However, when we started our work on this paper, HiSSE and GeoHiSSE could only deal with a small number of hidden states and had some inconsistencies in the computation of conditional probabilities (Herrera-Alsina et al. (2019). Hence, we worked with the SecSSE model (several examined and concealed states-dependent speciation and extinction (Herrera-Alsina et al. (2019), which combines the features of HiSSE with the MuSSE model for multiple states (Fitzjohn 2012), solves the limitations of previous SSE models, and can consider trait changes (here changes in biogeographical state) during speciation, as in the GeoSSE and ClaSSE (Goldberg and Igić 2012) models (see e.g. Aduse-Poku et al. 2021). The HiSSE package may now have the same features, but the SecSSE model is more familiar to us, as some of us developed it, which also allows us to optimize its performance.

The inferences that can be made with SSE models heavily rely on the quality of the data, but few biological groups have well-resolved phylogenies and complete trait-datasets. One such group is the Caninae subfamily, which have a well-resolved tree (Porto et al. 2019) and an incredibly detailed fossil record (Wang et al. 2004; Wang and Tedford 2008; Tedford et al. 2009). The rich and well-identified records of extinct Caninae species offer a unique opportunity to study the processes and mechanisms of their worldwide diversification when lineages reached new continents. Here, we present an extension of the SecSSE model (Herrera-Alsina et al. 2019; Aduse-Poku et al. 2021) to complete phylogenies (with all extinct species). We opted not to incorporate the birth-death-fossilization model (Stadler et al. 2018) because this would make the computations more complex than necessary for (nearly) complete phylogenies. We apply this new version to a complete tree of Caninae to study the effects of biogeographic states on canid rates of diversification and dispersal. We expected to find different speciation rates in South America and Africa due to the very distinct environments that both continents had in comparison with North America (Zachos et al. 2001; Strömberg 2011; Zhang et al. 2014), and also due to the absence of competitors in South America (Wang and Tedford 2008), which could be indicative of ecological opportunity. Furthermore, we expected extinction rates to be higher in Eurasia than elsewhere due to encounters with other groups of carnivores such as Felidae (Wang and Tedford 2008; Pires et al. 2015, 2017). However, our results do not support either scenario. Interestingly, our findings based on the complete tree for Caninae are very different from what we find if we only use extant species. This demonstrates the importance of using fossil information when available and being cautious in interpreting results when it is not available.

## 2. MATERIALS AND METHODS

### 2.1 Formulation

The SecSSE model assumes that speciation (*λ*_*ijk*_) and extinction rates (*μ*_*i*_) depend on the trait state *i* of the lineage, and in the case of speciation on the trait states *j* and *k* of the daughter branches. This trait is a combination of an observed trait and a hidden or concealed trait that can take discrete states which can change from state *I* to state *j* with a rate *q*_*ij*_. During speciation, the daughter species usually inherit the trait from the parent species, but SecSSE also allows the daughter species to have different trait states than the parent, and different from one another. In our implementation of SecSSE the trait is the biogeographic state, which can take the following values: eight states that we observe in the extant canids (North America (NAM), South America (SAM), Eurasia (EUR), Africa (AFR), NAM + SAM, NAM + EUR, EUR + AFR, and NAM + EUR + AFR), and seven more states which are currently unobserved but are natural alternatives to the eight observed ones (AFR + SAM, NAM + SAM + AFR, EUR + SAM, NAM + EUR + SAM, NAM + AFR, EUR + AFR + SAM, and EUR + NAM + SAM + AFR) (Figure 1A). Speciation in a species that is present in only one continent is always sympatric, which means that daughter species inherit the parent’s state. Speciation in a species that is present in multiple continents is always allopatric (vicariant), which means that the daughter species each inherit part of the range of the parent. Species can disperse to other continents thereby extending their ranges, and they can contract their ranges by going extinct in a single continent. Note that species in states comprising multiple continents cannot become extinct in a single step. They first need to contract their range until they are present in only one of the continents.

**Figure 1.**
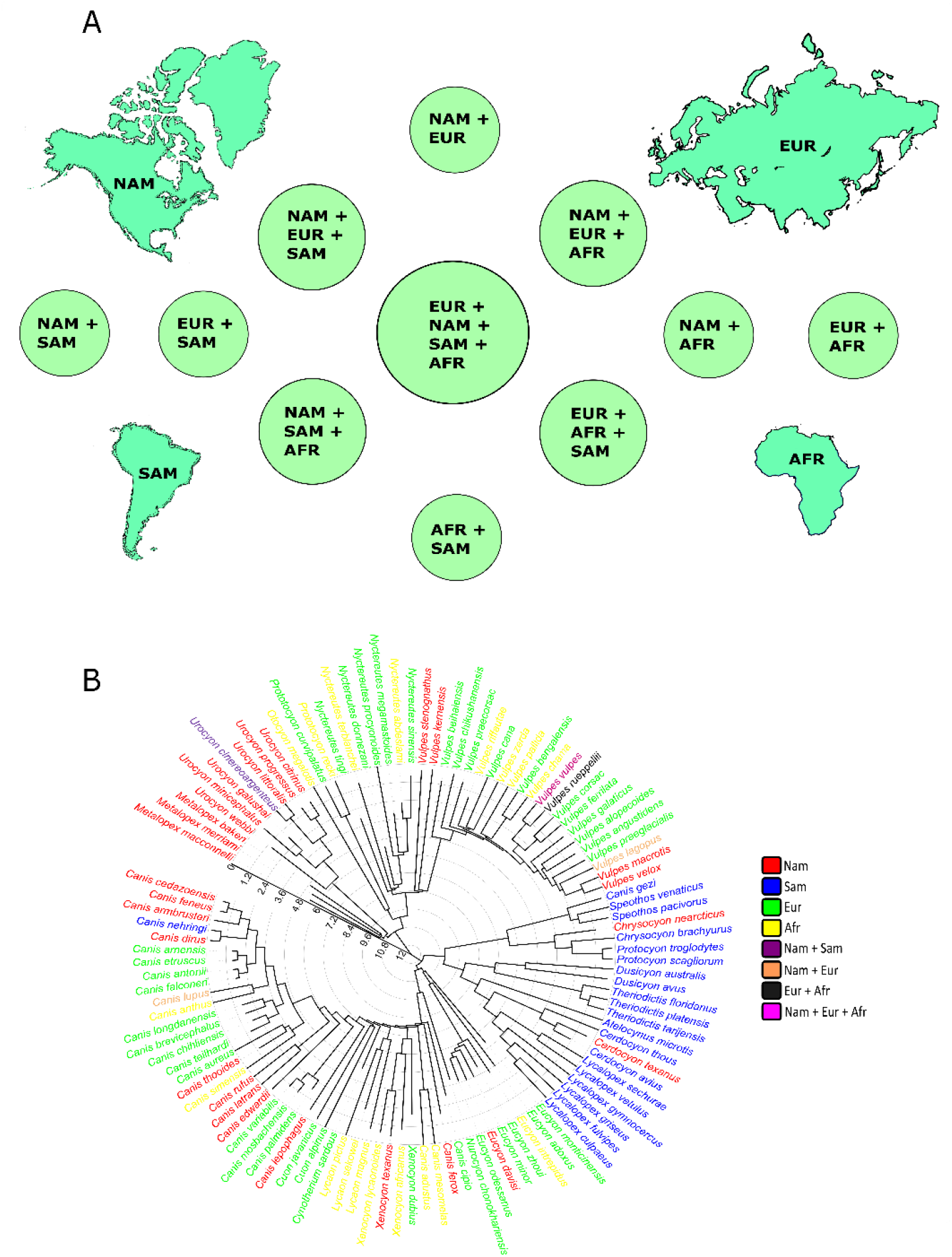
Trait states and phylogenetic tree used here; A) the four single states and their 11 combinations that we used to classify Caninae distribution patterns; B) the complete phylogenetic tree used here containing 111 species (75 extinct and 36 extant). The tips of the tree are colored based on their distributions.

Because the available version of the R package SecSSE only considered reconstructed trees without extinct species, we extended the package so it can be applied to complete trees (with all extinct species). For reconstructed trees, Goldberg and Igić (2012) provided equations to compute the likelihood *D*_*i*_(*t*) of the phylogeny subtending from a lineage at time *t* and the trait states at the present given trait state *i* at time *t*. By computing these probabilities backward in time (from tips to crown), and combining probabilities at the nodes, one can obtain the likelihood of the entire tree given the trait state at the crown. The equations also involve the extinction probability *E*_*i*_(*t*) which is the probability of the lineage at time *t* not having any descendants at the present. These formulas have been implemented in various packages including SecSSE. However, the mathematical formulation for complete trees is in fact much simpler as we no longer need to track the extinction probability because we observe all extinctions in the complete tree (but note that we still need to condition on survival of the process). The equation for *D*_*i*_(*t*) becomes:

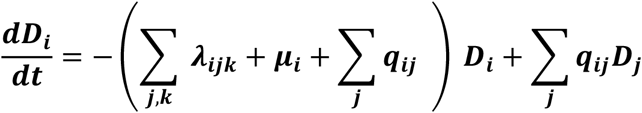

where we use, for the tree tips at the present, the initial condition *D*_*i*_ = 0 when the species is in state *i* at the tip and 0 otherwise, and for the extinct species *D*_*i*_ = *μ*_*i*_ when the species is in state *i* at the tip and 0 otherwise. At any node in the complete tree, the *D*_*i*_ of the daughter branches are used to compute the *D*_*i*_ of the parental branch:

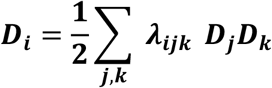

### 2.2 Phylogenetic tree and distribution data

The phylogeny produced by Porto et al. (2019) for the Caninae subfamily was used here as the basis for our tree. Their phylogeny, constructed with molecular and osteological data through Bayesian inference, has all the 36 extant canids in the world, plus the recently extinct (1876) *Dusicyon australis*. As we are measuring macroevolutionary extinction rates, *Dusicyon australis* was considered an extant species during our analyses because it was a human-induced extinction. We added all the other 74 extinct species described for Caninae following Wang and Tedford (2008), generating a complete tree for the subfamily. A literature review was performed in the digital paleobiology dataset Fossilworks (Alroy 1998) in order to define the most likely phylogenetic positions that each extinct species would take compared to the extant species in the phylogeny form Porto el al. (2019). Thus, we added the 74 extinct species in the phylogeny based on the age that was presented for each lineage in Fossilworks. This resulted in the phylogeny used in this study with 111 species (Figure 1B) (Table S1).

The complete Caninae tree was time-calibrated using the fossil information of the 74 extinct species obtained from Fossilworks, as well as the fossil ages used by Porto et al. (2019). This analysis was performed in *R* 4.0-2. (R Development Core Team 2020) using the *chronos* function of the ape 5.4-1 package (Paradis and Schliep 2019).

We divided the world into 15 biogeographic areas (trait states) (Figure 1A). We categorized all canids into the 15 previously mentioned biogeographical areas based on the distribution information for the 111 species obtained from IUCN (2020) and from Fossilworks (Table S1).

### 2.3 SecSSE models

We considered several models differing in whether observed (biogeography - i.e., presence in a biogeographic area) or hidden traits affect speciation and/or extinction rates or not, and whether transition rates (in our case range expansion and contraction) differ or not. Each model has one or several hypotheses being tested (Supplementary Material and Appendix 1), and all models followed the diversification scheme presented in Figure S1. We fitted 45 state-dependent speciation and extinction models. Among these models, 22 are ETD (Examined Trait-Dependent) models, 22 are CTD (Concealed-Trait-Dependent) models with the same set up as the ETD models, and one is a CR (Constant-Rate) model. In ETD models, the diversification rates depend on the trait of interest, i.e., the geographic range. In CTD models, the diversification rates depend on a concealed trait that we are not analyzing. This is the same as the hidden state in HiSSE but we stick to the terminology in Herrera-Alsina (2019) and hence use concealed. In the CR model, rates are homogeneous across states. The full set of models and all their rate parameters are detailed in the Appendix 1. Note that we allowed the same number of concealed and observed states (15), implying that our models have 225 possible combinations of examined and concealed states. We assumed that concealed states do not undergo changes during speciation, as this would lead to overparametrizing the model (and hence could result in nonsensical parameter estimates).

Speciation (*λ*_*i*_) matrices were set to only accommodate dual inheritance or dual symmetric transition scenarios by sympatric and allopatric speciation, respectively. Dual inheritance scenarios reflect that during speciation the trait state from the ancestor is perfectly inherited by both daughter species which in our case occurs for sympatric speciation. Dual symmetric transitions reflect that daughter species can inherit any state; in our case they inherit non-overlapping parts of the parental range (allopatric speciation). All 45 models have the same speciation matrix setup, but they vary in the number of different speciation parameters estimated Appendix 1. The various models consider 1) distinct rates for sympatric and allopatric speciation; 2) distinct rates for sympatric and allopatric speciation and a third rate for speciation in South America to study whether the endemic clade on this continent has a different diversification than the other continents; 3) a model with five speciation rates, two for sympatric and allopatric speciation in the new world (NAM and SAM and NAM+SAM), two for sympatric and allopatric speciation in the old world (EUR and AFR and EUR+AFR), and one for allopatric speciation among new and old world (for EUR+NAM and EUR+NAM+AFR). We refer to the Supplementary Material and to the Appendix 1 for a complete description of the hypothesis each model tests.

Extinction rates (*μ*_*i*_) were set for extinctions in the single-area states (NAM, SAM, EUR, and AFR). For the other 11 states (multiple-area states) extinction rates were fixed to zero; species in these states first have to contract their ranges to a single-area state before they can become extinct. We considered models with a distinct rate for one of the four single-area states while the other three had the same rate. Variations of this model were created for each one of the four single-area states to test if extinction was higher or lower in any of them compared to the others. Another model had four different rates for each one of the single-area states. We also had a model with two distinct extinction rates, one for old world states and another for new world states.

For the transition rates (*q*_*ij*_), we set constrained matrices for all the models, meaning that there are rules prohibiting some transitions. Transition rates from multiple-area states to single-area states are in fact extinctions because this transition means a local extinction in one of the areas that form the multiple-area state, and hence have the same rate *μ*_*i*_. The models considered here are: 1) distinct transition rates from old world to new world than from new world to old world; 2) distinct rates among states within new and old world; 3) rates for each of the possible transitions among areas; 4) and several models with two rates, one for one of the states and another rate for all others. Concealed states (CTD models) follow the same settings of *λ*_*i*_, *μ*_*i*_, and *q*_*ij*_ matrices used in the ETD models.

To run the models, we used the *cla_secsse_ml* function from the extended SecSSE package 3.0.0. We used Akaike Information Criterion (AIC) weights to compare CR, ETD and CTD models using the same data set (Akaike, 1973); these are AIC values that are normalized to add up to 1.

We used random initial values, drawn from a uniform distribution from 0 to 1. To lower chances of ending up in local optima, we set each optimization in the optimize function from the DDD 4.3 package (Etienne et al. 2016) to have 5 cycles (i.e., optimization starts again from the optimal parameters), and we repeated this for 10 different random initial parameter sets.

To test how much the interpretation of the evolutionary history of a group can change depending on the presence or absence of extinct species in phylogenies, we applied the 45 models not only to the complete phylogeny of Caninae (111 species), but also to the extant-species tree + one recently extinct (37 species) from Porto et al. (2019). Thus, the total number of analyses doubled to 90.

## 3. RESULTS

For the complete tree, we found that a CTD model had the highest support (model 37 – AIC weight = 0.737 – Figure 2A), in which rates of diversification are allowed to vary across lineages but independently of the areas (Table 1 - see Table S2 for the complete model comparison). Model 37 has three distinct speciation rates, which were estimated as *λ*_1_ = 0.045, *λ*_2_ = 3.13E-06, and *λ*_3_ = 0.811. This model has one *μ* for extinctions in all states (*μ*_1_ = 0.493). Although we found no effect of biogeographic areas on rates of diversification, we found substantial differences in rates of area expansion (*q*_*1*_ = 0.075, *q*_*2*_ = 0.013, *q*_*3*_ = 6.15E-16) and contraction (*μ*_1_ = 0.493) (Table S3).

**Table 1.**
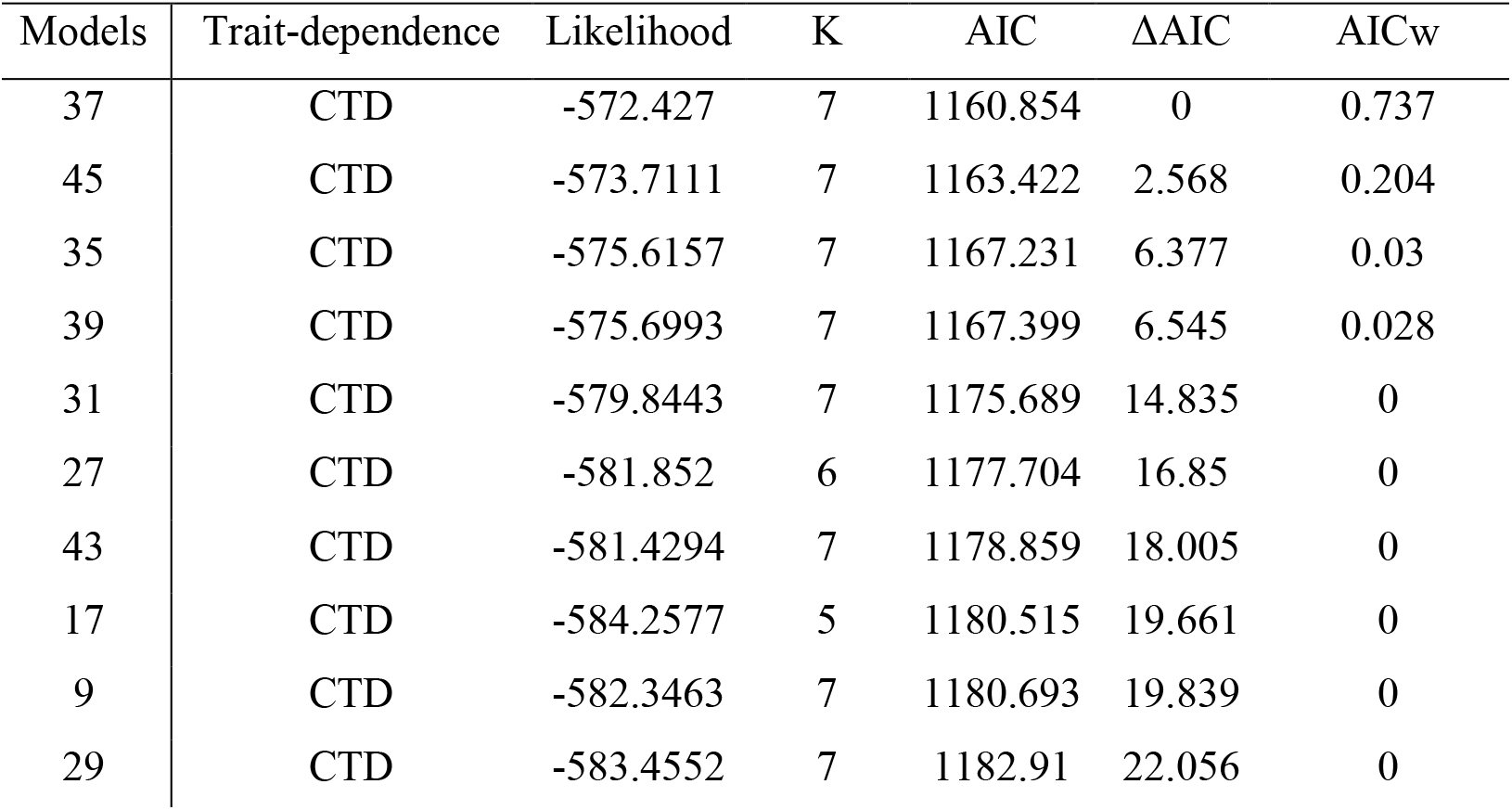
Model comparison based on AIC for the 10 best performing models tested with the complete tree. Models differ in what state changes are allowed, and whether state changes occur during speciation. CR = Constant Rates; CTD = Concealed trait-dependent; ETD = Examined trait-dependent; K = number of free parameters. The highest supported model (model 37) is the CTD version of the ETD model 36, which has three speciation rates and one extinction rate, plus three transition rates. The area contraction rate is the same as *μ*_1_.

**Figure 2.**
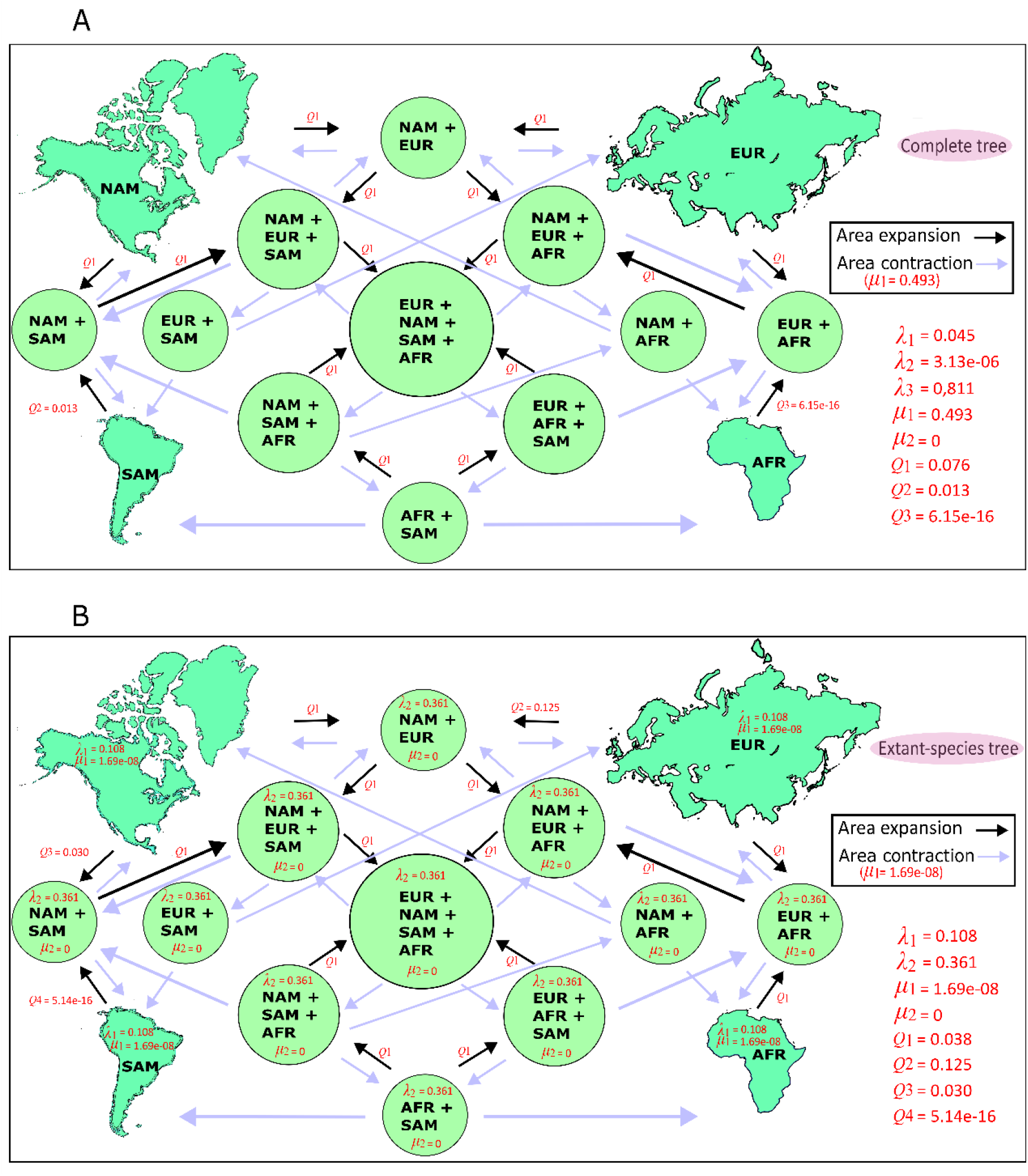
Estimates of rates of speciation (*λ*), extinction (*μ*) and transition across states (*q*_*ij*_) for the best supported models (see Tables S2 – S5). A) In the scenario where we assessed the complete tree, CTD models had better results, which suggests that other factors than biogeographic areas cause variation in diversification rates. B) For the extant-species tree, ETD models performed better, suggesting that the geographic distribution partly drives Caninae diversification.

There was no ETD model among the 10 best models (Table 1).

Among the 45 models we compared for the extant-species tree, we found the highest support for an ETD model (model 30 – AIC weight = 0.365 – Figure 2B) (See Table S4 for the model comparison). This model assumes distinct speciation rates for sympatric and allopatric speciation, and explores whether Eurasian lineages expanded more their distributions to North America or North American lineages expanded more to Eurasia (if there is a higher transition to N. America from Eurasia than the other way around, which could suggest that Felidae imposed an incumbent effect on Canids throughout Beringia (Pires et al., 2015)). The model also explores how strong the Panama bridge was as a filter for N. American lineages and S. American lineages that tried to cross it. The estimated speciation rates were *λ*_1_ = 0.109, and *λ*_2_ = 0.361 (Table S5). Model 30 had only one *μ* estimated that represented extinctions in all the single-area states and also area contraction (*μ*_1_ = 1.69E-08). This model assumes four free transition rates: *q*_*2*_ = 0.126 for the transition from Eurasia to N. America; *q*_*3*_ = 0.031 for the transition from N. America to S. America; *q*_*4*_ = 5.14E-16 for transitions from S. America to N. America; and *q*_*1*_ = 0.038 for transitions among all other combinations of areas.

The second-best model for the extant-species tree was also an ETD model, i.e., model 22, with a ΔAIC = 0.951 and an AIC weight = 0.227 (Table S4). This model has a very similar setup as model 30, only differing in the *q* matrix. Model 22 does not test differential transition rates from Eurasia to N. America, like model 30 does, but assumes the same speciation rates for both biogeographic areas.

Support for ETD models was generally high, regardless of the setup of each model, summing up to an AIC weight of 0.867 (note that our set of models is balanced; for every ETD model there is a corresponding CTD model).

Models 40, 42 and 44, that were set up to test whether the centers of origin of the three major clades of Caninae (South America, North America, and Africa) have different speciation rates than the other areas, did not perform much better than models that assumed these rates were identical, such as models 32 and 36. In addition, models with area-dependent extinction did not perform better than models with area-dependent speciation.

Rates of allopatric speciation have a very wide range of values, from 0.08 to 0.58, but most of the estimations are around 0.27 (Figure 3A), being the highest lambda parameter found among our extant-species tree models, and more than two times higher than sympatric speciation. Among the single-area states, on average, S. America presented the highest lambda (0.18).

**Figure 3.**
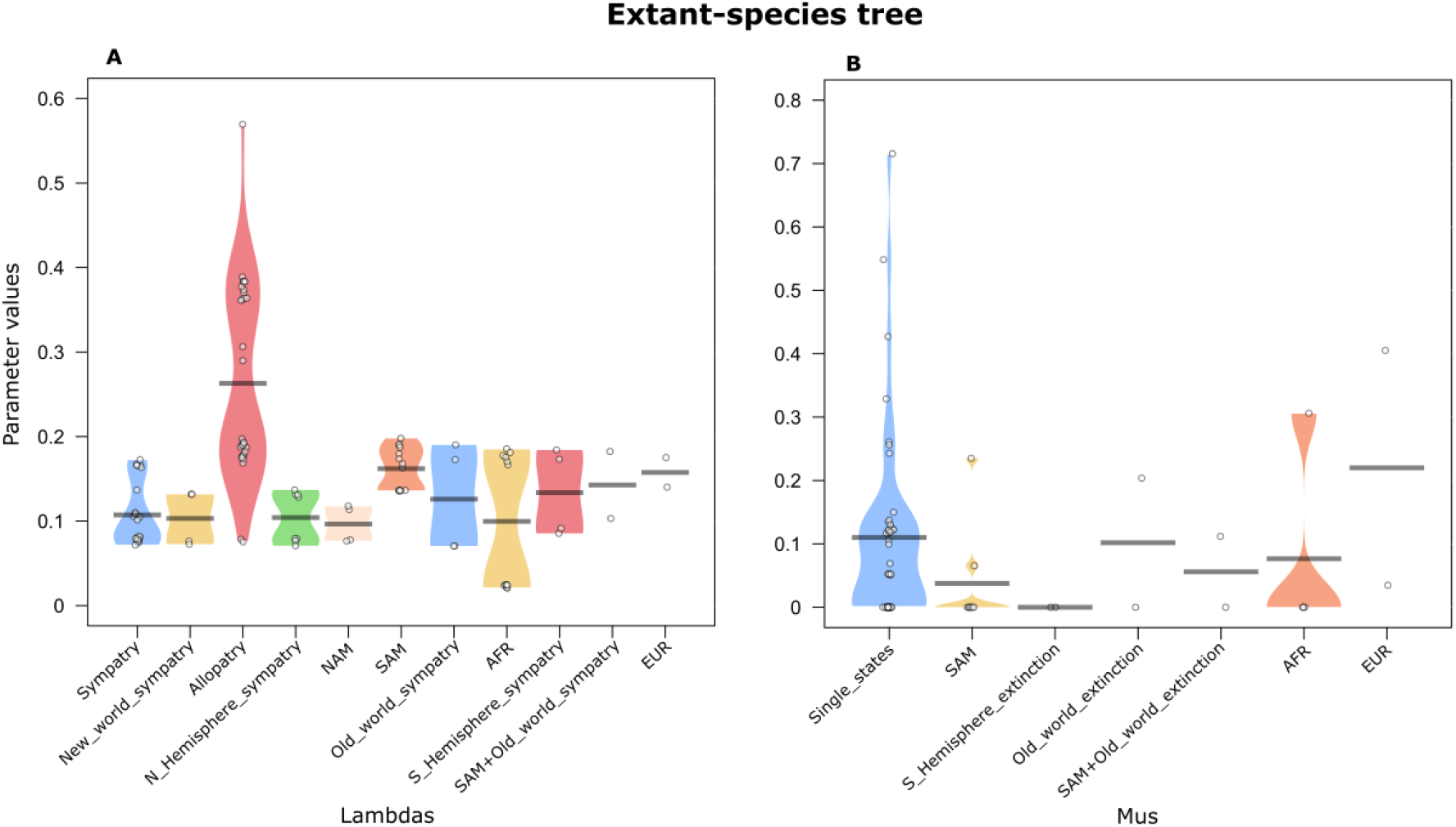
Comparison among all the diversification rates estimated across the 45 models tested here for the extant-species tree. On the left (A) the speciation rates, and on the right (B) the extinction rates. “EUR”, “AFR”, “NAM” and “SAM” mean a separate *λ* or *μ* for each of these continents; “allopatry” and “sympatry” mean a separate *λ* for these speciation events; and “single-states” means one single *λ* or *μ* for each state.

## 4. DISCUSSION

Our analyses, carried out separately for the reconstructed and the complete phylogeny, show that taking into account fossil information in SecSSE models substantially changes the interpretation of how biogeographic areas influence diversification rates of Caninae lineages. In the scenario with only extant species, we found better performance of ETD models, suggesting that the distinct diversification patterns that we see on the phylogeny of extant canids are related to geographic areas. By contrast, when extinct species were incorporated into the phylogeny, we found that CTD models performed much better than ETD models. This indicates that the rates of diversification for all known Caninae species cannot be explained by continent-related biogeographic events.

The best scenario for the extant-species tree, model 30, shows very appealing results as Eurasian lineages are more unlikely to expand their distributions into North America than vice versa, supporting the idea of an incumbency effect in Beringia (Rosenzweig and Mccord 1991) due to the presence of other carnivores, as Felidae, hindering the passage of competing lineages from N. America, which was also proposed by Pires et al. (2015). Furthermore, model 30 indicates that South American canids found it more challenging to cross the Panama bridge compared to North American canids. In the literature, one could find an explanation for this observed pattern as significant diversification events occurred in S. America, giving rise to numerous endemic species that became highly specialized for life within the forested environment of the continent (Porto et al., 2023). One could argue that such specialization made it challenging for South American canids to transition back to open habitats such as those in North America at that moment. However, as tempting as this idea is, the extinction rates for all continents are very low, and we know from the observation of 75 extinct species, that this value cannot be correct. Therefore, the low extinction rate for the extant-species phylogeny together with the CTD model being chosen for the complete tree practically disqualifies any interpretation we can get from the extant-species tree analyses. Thus, our interpretations of the evolution of whole groups can radically change when fossil information is incorporated.

With respect to the models with the complete tree, no effect of biogeographic areas on rates of diversification was found, but there were differences in transition rates. According to the best supported model, area contraction was substantially faster than area expansion. Model 37 also suggests that it was difficult for species to disperse from Africa to Eurasia. Models 45 and 39 also assess this scenario with a differential transition rate to dispersion events from Africa to Eurasia, also exhibiting low *q* rates compared to the rest of the states. In addition to our best model, models 45 and 39 rank as the second and fourth best models by AIC. Thus, the pattern of limited expansion out of Africa persists in models incorporating this hypothesis, even with varying parameter structures. The incumbency effect can be a good explanation for these low values of area expansion. To be able to disperse into new continents, Caninae lineages needed to pass through very narrow land areas such as portions of land that connect Africa with Eurasia. These places probably had the presence of other carnivores that would hinder the passage of canids. For example, it was suggested by Silvestro et al. (2015) that Felidae might have contributed to increase the extinction rate in North American canids.

There are few possibilities why geographic areas are not important for the diversification of canids in our complete tree scenario. It is very unlikely that major events of dispersal around the world did not have a large impact on lineages (Porto et al. 2021), but maybe such dispersal events are more important at smaller scales, particularly in intercontinental connections. Here, phenomena such as incumbent effects imposed by other carnivores on canids could gain prominence, aligning with some of the leading models for both the complete tree and extant-species tree. Yet, at larger scales, additional intrinsic factors of the areas might intersect with our geographic trait, potentially introducing a concealed layer of complexity not fully captured by our models. For example, studies by Rolland et al. (2014) and van Els et al. (2021) highlight hidden variations within latitude and elevation gradients.

Furthermore, probably there is one or there are a few traits that these fossil species carry with them, and are essential for a complete understanding of Caninae, which turns out to be very compatible with the scenario we have here, as 67% of the species in our complete tree are extinct. This leads us to hypothesize that extinct species may have different factors determining their (absence of) diversification than extant species. Two traits present themselves as potential candidates: 1) diet variation, such as omnivory, can be a strategy for surviving based on resource availability (Ingram et al. 2009) and, if lineages shift to omnivorous habits during moments of environmental perturbation, this may lead to low diversification (Van Valkenburgh et al. 2004). This could explain diversification of the South American clade, which contains several species with omnivorous diets; 2) body size is generally believed to contribute to diversification, because smaller species of mammals tend to have higher speciation rates, while larger species tend to have larger extinction probabilities (Liow et al. 2008). Both traits can be tested through a SecSSE analysis.

In summary, our findings suggest that even though there is heterogeneity in rates of diversification for extant canids, the biogeographic area where the species occurs does not seem to drive this heterogeneity when we incorporate fossil species in our analyses. Thus, we highlight the effect that the inclusion of fossil information in our models has on our understanding about the evolution of Caninae. In addition, we propose that more complex models can help our understanding about evolutionary dynamics. As lineages disperse to new continents, other traits (e.g., diet and body size) may have played prominent roles in the evolution of species and, together with distribution patterns, could bring a more complete scenario about diversification events through time.

## Notes

### Competing Interest Statement

The authors have declared no competing interest.

