## Supplementary material for "From Fossils to Living Canids: Two Contrasting Perspectives on Biogeographic Diversification"

### DESCRIPTIONS OF THE MODELS (ETDS)

**Model 1 -** Constant-Rate model. This model has 3 free parameters to be estimated ($\lambda_1$, $\mu_1$ and $q_1$). Transitions from multiple-area states to single-area states were regarded as extinctions ($\mu_1$). In the CR model, all species have the same speciation rate regardless of their trait state (e.g., $\lambda_{1A} = \lambda_{1B} = \lambda_{1C}$), where the numbers refer to the observed and examined biogeographic state and the letter to the unknown hidden trait. For all models, the extinction rate for multiple-area states was fixed to zero ($\mu_2 = 0$), and transitions from multiple-area states to single-area states were regarded as extinctions ($\mu_1$).

**Model 2 -** This ETD model has 4 free parameters to be estimated ($\lambda_1$, $\lambda_2$, $\mu_1$ and $q_1$). Transitions from multiple-area states to single-area states were regarded as extinctions ($\mu_1$). In this ETD model, speciation rates are allowed to vary only for the examined states, and then only between sympatric and allopatric speciation. This model assumes that there is a difference between sympatric and allopatric speciation in canid diversification.

**Model 4 -** This ETD model has 4 free parameters to be estimated ($\lambda_1$, $\mu_1$, $q_1$ and $q_2$). It is a generalization of the CR model (model 1) with one more transition rate. This model tests if transitions among new and old world were lower or higher than transitions within old world and new world. Here we are interested if crossing the Bering land-bridge was more difficult than crossing to areas within new and old world. Some evidence suggests that crossing from North America to Eurasia was difficult for some lineages due the felines in Siberia (incumbency effect).

**Model 6 -** This ETD model is similar to model 2, but with one more transition rate (5 free parameters - $\lambda_1$, $\lambda_2$, $\mu_1$, $q_1$ and $q_2$). Here assume that there is a difference between sympatric and allopatric speciation in canid diversification, and we test if transitions among new and old world were lower or higher than transitions within old world and new world (we are interested if crossing the Bering land-bridge was more difficult than crossing to areas within new and old world). Some evidence suggests that crossing from North America to Eurasia was difficult for some lineages due the felines in Siberia (incumbency effect).

**Model 8 -** This ETD model has 7 free parameters to be estimated ($\lambda_1$, $\lambda_2$, $\lambda_3$, $\mu_1$, $\mu_3$, $q_1$ and $q_2$). Transitions from multiple-area states to single-area states were regarded as extinctions ($\mu_1$). This model tests for 1) a difference between sympatric and allopatric speciation in canid diversification; 2) distinct speciation and extinction rates in S. America.

**Model 10 -** This ETD model has 4 free parameters to be estimated ($\lambda_1$, $\lambda_2$, $\mu_1$ and $q_1$). This model tests fora distinct speciation rate of S. America compared to the rest of the world.

**Model 12 -** This ETD model has 4 free parameters to be estimated ($\lambda_1$, $\mu_1$, $\mu_3$ and $q_1$). This model assumes a distinct speciation rate for S. America compared to the rest of the world and a specific rate of extinction in S. America.

**Model 14 -** This ETD model has 4 free parameters to be estimated) ($\lambda_1$, $\mu_1$, $q_1$ and $q_2$). This model assumes distinct speciation and extinction rates for old and new world and a distinct transition rate to South America.

**Model 16 -** This ETD model has 5 free parameters to be estimated ($\lambda_1$, $\lambda_2$, $\lambda_3$, $\mu_1$ and $q_1$). This model tests for distinct speciation rates for sympatric speciation in new and old worlds, and a third rate for allopatric speciation. The justification for exploration of this model is that North American lineages of carnivores invading Eurasia underwent explosive radiations, whereas lineages invading North America maintained uniform diversification dynamics.

**Model 18 -** This ETD model has 6 free parameters to be estimated ($\lambda_1$, $\lambda_2$, $\mu_1$, $\mu_3$, $\mu_4$ and $q_1$). Here we test if extinction rates were higher in North America, because Pires et al. (2015) suggest that diversification rates of carnivores in this continent were lower than in Eurasia, which could be due to the entrance of felines from Eurasia hunting for and competing with canids.

**Model 20 -** This ETD model has 5 free parameters to be estimated ($\lambda_1$, $\mu_1$, $q_1$, $q_2$, and $q_3$). This model tests if South American lineages expanded more their distributions to North America or if North American lineages expanded more to the south. This explores how strong the Panama land-bridge was as a filter for the lineages in both regions.

**Model 22 -** This ETD model has 6 free parameters to be estimated ($\lambda_1$, $\lambda_2$, $\mu_1$, $q_1$, $q_2$, and $q_3$). This model assumes distinct speciation rates for sympatric and allopatric speciation, and explores if South American lineages expanded more their distributions to North America or if North American lineages expanded more to the south. This explores how strong the Panama land-bridge was as a filter for the lineages in regions.

**Model 24 -** This ETD model has 5 free parameters to be estimated ($\lambda_1$, $\lambda_2$, $\lambda_3$, $\mu_1$, and $q_1$). This model assumes the same speciation rate for sympatric and allopatric speciation, and two distinct speciation rates for Africa and S. America. Both regions seem to be hotspots of diversification – Africa (fox clade) and S. America (S. American clade).

**Model 26 -** This ETD model has 6 free parameters to be estimated ($\lambda_1$, $\lambda_2$, $\lambda_3$, $\lambda_4$, $\mu_1$, and $q_1$). This model assumes distinct speciation rates for sympatric and allopatric speciation, and distinct speciation rates for Africa and S. America. Both regions seem to be hotspots of diversification – Africa (fox clade) and S. America (S. American clade).

**Model 28 -** This ETD model has 7 free parameters to be estimated ($\lambda_1$, $\lambda_2$, $\lambda_3$, $\lambda_4$, $\lambda_5$, $\mu_1$, and $q_1$). This model assumes distinct speciation rates for sympatric and allopatric speciation, and distinct dynamics of sympatric speciation in 4 areas.

**Model 30 -** This ETD model has 7 free parameters to be estimated ($\lambda_1$, $\lambda_2$, $\mu_1$, $q_1$, $q_2$, $q_3$, and $q_4$). This model assumes distinct speciation rates for sympatric and allopatric speciation, and explores if Eurasian lineages expanded more their distributions to North America or North American lineages expanded more to Eurasia (if there is a higher transition to N. America from Eurasia than the other way around, this could suggest that Felidae really imposed an incumbent effect on Canids throughout Beringia). The model also explores how strong the Panama bridge was as a filter for N. American lineages and S. American lineages that tried to cross it.

**Model 32 -** This ETD model has 7 free parameters to be estimated ($\lambda_1$, $\lambda_2$, $\lambda_3$, $\lambda_4$, $\mu_1$, $\mu_3$, and $q_1$). This model assumes distinct speciation rates for sympatric and allopatric speciation, and assumes one speciation rate for S. America and another for Africa. Both regions seem to be hotspots of diversification – Africa (fox clade) and S. America (S. American clade).

**Model 34 -** This ETD model has 7 free parameters to be estimated ($\lambda_1$, $\lambda_2$, $\lambda_3$, $\mu_1$, $\mu_3$, $\mu_4$, and $q_1$). This model assumes distinct speciation rates for sympatric and allopatric speciation, and the same speciation rate for S. America and Africa. Both regions seem to be hotspots of diversification – Africa (fox clade) and S. America (S. American clade).

**Model 36 -** This ETD model has 7 free parameters to be estimated ($\lambda_1$, $\lambda_2$, $\lambda_3$, $\mu_1$, $q_1$, $q_2$, and $q_3$). This model assumes distinct speciation rates for sympatric and allopatric speciation, and the same speciation rate for S. America and Africa, and assumes three distinct transitions, one for transitions from S. America, another for transitions from Africa, and one for the rest of the continents. Both regions seem to be hotspots of diversification – Africa (fox clade) and S. America (S. American clade).

**Model 38 -** This ETD model has 7 free parameters to be estimated ($\lambda_1$, $\lambda_2$, $\lambda_3$, $\mu_1$, $q_1$, $q_2$, and $q_3$). This model assumes the same speciation rate for sympatric and allopatric speciation, and two distinct speciation rates for S. America and Africa, and assumes three transitions, one for transitions from S. America, another for transitions from Africa, and one for the rest of the continents. Both regions seem to be hotspots of diversification – Africa (fox clade) and S. America (S. American clade).

**Model 40 -** This ETD model has 6 free parameters to be estimated ($\lambda_1$, $\lambda_2$, $\lambda_3$, $\mu_1$, $\mu_3$, and $q_1$). This model assumes distinct speciation rates for sympatric speciation in new and old worlds, and another rate for allopatric speciation, and two distinct extinction rates for new and old worlds.

**Model 42 -** This ETD model has 7 free parameters to be estimated ($\lambda_1$, $\lambda_2$, $\lambda_3$, $\mu_1$, $\mu_3$, $q_1$, and $q_2$). This model assumes distinct speciation rates for sympatric and for allopatric speciation, a distinct rate of speciation for N. America, two distinct extinction rates, one for N. America and the other for the rest of the single-area states, and one transition rate from N. America and another for transitions between the other regions. We explore distinct dynamics for N. America, because canids originated here, so the diversification may have been different.

**Model 44 -** This ETD model has 7 free parameters to be estimated ($\lambda_1$, $\lambda_2$, $\lambda_3$, $\mu_1$, $\mu_3$, $q_1$, and $q_2$). This model assumes distinct speciation rates for sympatric and for allopatric speciation, a distinct rate of speciation for Africa, two distinct extinction rates, one for Africa and the other for the rest of the single-area states, and one transition rate from Africa and another for transitions between the rest of the regions. We explore distinct dynamics for N. America, because canids originated here, so the diversification may have been different.

### TABLE LIST

**Table S1**. List of the 111 species of Caninae included in our study with the distribution areas that they belong based on our 15 biogeographical regions used here. Species marked with (*) are the extinct canids include in the tree of Porto et al. (2019).

| Species | State |
|---|---|
| *Canis lupus* | NAM+EUR |
| *Canis anthus* | AFR |
| *Canis aureus* | EUR |
| *Canis simensis* | AFR |
| *Canis rufus* | NAM |
| *Canis latrans* | NAM |
| *Cuon alpinus* | EUR |
| *Lycaon pictus* | AFR |
| *Canis adustus* | AFR |
| *Canis mesomelas* | AFR |
| *Lycalopex vetulus* | SAM |
| *Lycalopex sechurae* | SAM |
| *Lycalopex gymnocercus* | SAM |
| *Lycalopex culpaeus* | SAM |
| *Lycalopex fulvipes* | SAM |
| *Lycalopex griseus* | SAM |
| *Cerdocyon thous* | SAM |
| *Atelocynus microtis* | SAM |
| *Dusicyon australis* | SAM |
| *Chrysocyon brachyurus* | SAM |
| *Speothos venaticus* | SAM |
| *Vulpes rueppellii* | EUR+AFR |
| *Vulpes vulpes* | NAM+EUR+AFR |
| *Vulpes ferrilata* | EUR |
| *Vulpes corsac* | EUR |
| *Vulpes velox* | NAM |
| *Vulpes macrotis* | NAM |
| *Vulpes lagopus* | NAM+EUR |
| *Vulpes chama* | AFR |
| *Vulpes bengalensis* | EUR |
| *Vulpes pallida* | AFR |
| *Vulpes zerda* | AFR |
| *Vulpes cana* | EUR |
| *Nyctereutes procyonoides* | EUR |
| *Otocyon megalotis* | AFR |
| *Urocyon littoralis* | NAM |
| *Urocyon cinereoargenteus* | NAM |
| *Canis dirus** | NAM+SAM |
| *Canis armbrusteri** | NAM |

| | |
|---|---|
| *Cuon javanicus** | EUR |
| *Canis ferox** | NAM |
| *Canis edwardii** | NAM |
| *Lycaon magnus** | AFR |
| *Canis lepophagus** | NAM |
| *Vulpes riffautae** | AFR |
| *Cerdocyon avius** | NAM+SAM |
| *Chrysocyon nearcticus** | NAM |
| *Dusicyon avus** | SAM |
| *Canis nehringi** | SAM |
| *Protocyon troglodytes** | SAM |
| *Protocyon scagliorum** | SAM |
| *Nyctereutes donnezani** | EUR |
| *Nyctereutes megamastoides** | EUR |
| *Speothos pacivorus** | SAM |
| *Vulpes stenognathus** | NAM |
| *Vulpes kernensis** | NAM |
| *Vulpes alopecoides** | EUR |
| *Vulpes praecorsac** | EUR |
| *Vulpes angustidens** | EUR |
| *Vulpes galaticus** | EUR |
| *Vulpes chikushanensis** | EUR |
| *Vulpes beihaiensis** | EUR |
| *Vulpes praeglacialis** | EUR |
| *Metalopex bakeri** | NAM |
| *Metalopex merriami** | NAM |
| *Metalopex macconnelli** | NAM |
| *Urocyon citrinus** | NAM |
| *Urocyon progressus** | NAM |
| *Urocyon galushai** | NAM |
| *Urocyon minicephalus** | NAM |
| *Prototocyon recki** | AFR |
| *Prototocyon curvipalatus** | EUR |
| *Nyctereutes tingi** | EUR |
| *Nyctereutes sinensis** | EUR |
| *Nyctereutes abdeslami** | AFR |
| *Nyctereutes terblanchei** | AFR |
| *Theriodictis floridanus** | NAM+SAM |
| *Theriodictis tarijensis** | SAM |
| *Theriodictis platensis** | SAM |
| *Eucyon intrepidus** | AFR |
| *Eucyon minor** | EUR |
| *Eucyon davisi** | NAM+EUR |
| *Eucyon zhoui** | EUR |

| | |
|---|---|
| *Eucyon monticinensis** | EUR |
| *Eucyon adoxus** | EUR |
| *Eucyon odessanus** | EUR |
| *Nurocyon chonokhariensis** | EUR |
| *Canis cipio** | EUR |
| *Canis etruscus** | EUR |
| *Canis falconeri** | EUR |
| *Canis arnensis** | EUR |
| *Canis antonii** | EUR |
| *Canis chihliensis** | EUR |
| *Canis palmidens** | EUR |
| *Canis variabilis** | EUR |
| *Canis teilhardi** | EUR |
| *Canis longdanensis** | EUR |
| *Canis brevicephalus** | EUR |
| *Cynotherium sardous** | EUR |
| *Canis cedazoensis** | NAM |
| *Canis gezi** | SAM |
| *Canis mosbachensis** | EUR |
| *Xenocyon africanus** | AFR |
| *Lycaon sekowei** | AFR |
| *Xenocyon dubius** | EUR |
| *Xenocyon texanus** | NAM |
| *Xenocyon lycaonoides** | NAM+EUR+AFR |
| *Cerdocyon texanus** | NAM |
| *Canis feneus** | NAM |
| *Canis thöoides** | NAM |
| *Urocyon webbi** | NAM |

**Table S2**. Model comparison based on AIC for the models tested with the complete tree. Models differ in what state changes are allowed, and whether state changes occur during speciation. CR = Constant Rates; CTD = Concealed trait-dependent; ETD = Examined trait-dependent; K = number of free parameters.

| Model | Trait-dependence | loglikelihood | K | AIC | ΔAIC | AICw |
|---|---|---|---|---|---|---|
| 37 | CTD | -572.427 | 7 | 1160.854 | 0 | 0.737 |
| 45 | CTD | -573.7111 | 7 | 1163.422 | 2.568 | 0.204 |
| 35 | CTD | -575.6157 | 7 | 1167.231 | 6.377 | 0.03 |
| 39 | CTD | -575.6993 | 7 | 1167.399 | 6.545 | 0.028 |
| 31 | CTD | -579.8443 | 7 | 1175.689 | 14.835 | 0 |
| 27 | CTD | -581.852 | 6 | 1177.704 | 16.85 | 0 |
| 43 | CTD | -581.4294 | 7 | 1178.859 | 18.005 | 0 |
| 17 | CTD | -584.2577 | 5 | 1180.515 | 19.661 | 0 |
| 9 | CTD | -582.3463 | 7 | 1180.693 | 19.839 | 0 |
| 29 | CTD | -583.4552 | 7 | 1182.91 | 22.056 | 0 |
| 33 | CTD | -583.9897 | 7 | 1183.979 | 23.125 | 0 |
| 25 | CTD | -586.8822 | 5 | 1185.764 | 24.91 | 0 |
| 7 | CTD | -588.4322 | 5 | 1188.864 | 28.01 | 0 |
| 41 | CTD | -587.5817 | 6 | 1189.163 | 28.309 | 0 |
| 23 | CTD | -587.9581 | 6 | 1189.916 | 29.062 | 0 |
| 5 | CTD | -590.0413 | 4 | 1190.083 | 29.229 | 0 |
| 11 | CTD | -590.7268 | 4 | 1191.454 | 30.6 | 0 |
| 19 | CTD | -589.1349 | 6 | 1192.27 | 31.416 | 0 |
| 21 | CTD | -592.0723 | 5 | 1196.145 | 35.291 | 0 |
| 15 | CTD | -594.4474 | 4 | 1198.895 | 38.041 | 0 |
| 3 | CTD | -594.5538 | 4 | 1199.108 | 38.254 | 0 |
| 13 | CTD | -596.5876 | 4 | 1203.175 | 42.321 | 0 |
| 28 | ETD | -594.1396 | 7 | 1204.279 | 43.425 | 0 |
| 38 | ETD | -595.167 | 7 | 1206.334 | 45.48 | 0 |
| 26 | ETD | -596.3111 | 6 | 1206.622 | 45.768 | 0 |
| 24 | ETD | -597.5327 | 5 | 1207.065 | 46.211 | 0 |
| 32 | ETD | -595.7353 | 7 | 1207.471 | 46.617 | 0 |
| 44 | ETD | -595.8706 | 7 | 1207.741 | 46.887 | 0 |
| 30 | ETD | -595.9647 | 7 | 1207.929 | 47.075 | 0 |
| 42 | ETD | -596.8718 | 7 | 1209.744 | 48.89 | 0 |
| 36 | ETD | -597.4017 | 7 | 1210.803 | 49.949 | 0 |
| 34 | ETD | -597.4537 | 7 | 1210.907 | 50.053 | 0 |
| 4 | ETD | -600.6105 | 4 | 1211.221 | 50.367 | 0 |
| 6 | ETD | -600.0606 | 5 | 1212.121 | 51.267 | 0 |
| 20 | ETD | -601.0596 | 5 | 1214.119 | 53.265 | 0 |
| 22 | ETD | -600.6667 | 6 | 1215.333 | 54.479 | 0 |
| 12 | ETD | -602.8305 | 4 | 1215.661 | 54.807 | 0 |
| 14 | ETD | -602.9298 | 4 | 1215.86 | 55.006 | 0 |
| 1 | CR | -604.3386 | 3 | 1216.677 | 55.823 | 0 |
| 18 | ETD | -601.3754 | 6 | 1216.751 | 55.897 | 0 |
| 2 | ETD | -604.0787 | 4 | 1218.157 | 57.303 | 0 |
| 8 | ETD | -601.1096 | 7 | 1218.219 | 57.365 | 0 |
| 16 | ETD | -603.238 | 5 | 1218.476 | 57.622 | 0 |
| 10 | ETD | -604.2811 | 4 | 1218.562 | 57.708 | 0 |
| 40 | ETD | -603.1604 | 6 | 1220.321 | 59.467 | 0 |

**Table S3.** Estimates of all the parameters used in each model for the complete tree. Models are ordered from best fitted model to the worst.

| Model | $\lambda_1$ | $\lambda_2$ | $\lambda_3$ | $\lambda_4$ | $\lambda_5$ | $\mu_1$ | $\mu_2$ | $\mu_3$ | $\mu_4$ | $q_1$ | $q_2$ | $q_3$ | $q_4$ |
|---|---|---|---|---|---|---|---|---|---|---|---|---|---|
| 37 | 0.046 | 3.13E-06 | 0.812 | - | - | 0.493 | 0 | - | - | 0.0766 | 0.0131 | 6.15E-16 | - |
| 45 | 0.022 | 1.03E-07 | 0.495 | - | - | 1.253 | 0 | 4.96E-07 | - | 0.0803 | 0.0238 | - | - |
| 35 | 5.89E-14 | 0.482 | 6.74E-07 | - | - | 1.745 | 0 | 0.088 | 6.66E-09 | 0.0724 | - | - | - |
| 39 | 0.035 | 3.17E-08 | 0.635 | - | - | 0.474 | 0 | - | - | 0.0752 | 0.0128 | 4.58E-15 | - |
| 31 | 0.035 | 0.685 | - | - | - | 0.610 | 0 | - | - | 3.59E-08 | 0.1113 | 0.0535 | 0.0233 |
| 27 | 0.044 | 0.019 | 0.096 | 0.832 | - | 0.464 | 0 | - | - | 0.0676 | - | - | - |
| 43 | 0.012 | 0.040 | 0.685 | - | - | 1.117 | 0 | 0.550 | - | 0.1039 | 0.0156 | - | - |
| 17 | 0.039 | 3.50E-14 | 0.826 | - | - | 0.534 | 0 | - | - | 0.0629 | - | - | - |
| 9 | 0.019 | 0.727 | 0.059 | - | - | 1.396 | 0 | 4.21E-07 | - | 0.1213 | 0.0215 | - | - |
| 29 | 0.058 | 4.95E-08 | 0.011 | 0.917 | 0.121 | 0.396 | 0 | - | - | 0.0652 | - | - | - |
| 33 | 0.022 | 0.104 | 0.140 | 0.990 | - | 1.322 | 0 | 0.225 | - | 0.0792 | - | - | - |
| 25 | 0.046 | 1.14E-07 | 0.787 | - | - | 0.399 | 0 | - | - | 0.0652 | - | - | - |
| 7 | 0.040 | 0.697 | - | - | - | 0.555 | 0 | - | - | 0.0849 | 3.08E-16 | - | - |
| 41 | 0.038 | 7.36E-07 | 0.911 | - | - | 0.451 | 0 | 0.914 | - | 0.0788 | - | - | - |
| 23 | 0.033 | 0.630 | - | - | - | 0.368 | 0 | - | - | 0.0814 | 0.0232 | 0.0212 | - |
| 5 | 0.105 | - | - | - | - | 0.462 | 0 | - | - | 0.0746 | 3.10E-16 | - | - |
| 11 | 0.038 | 0.727 | - | - | - | 0.400 | 0 | - | - | 0.0627 | - | - | - |
| 19 | 0.004 | 0.650 | - | - | - | 0.425 | 0 | 0.736 | 1.263 | 0.0809 | - | - | - |
| 21 | 0.107 | - | - | - | - | 0.537 | 0 | - | - | 0.0882 | 0.0247 | 0.0225 | - |
| 15 | 0.105 | - | - | - | - | 0.393 | 0 | - | - | 0.0230 | - | - | - |
| 3 | 0.032 | 0.629 | - | - | - | 0.357 | 0 | - | - | 0.0668 | - | - | - |
| 13 | 0.106 | - | - | - | - | 1.080 | 0 | 0.222 | - | 0.0917 | - | - | - |
| 28 | 0.184 | 0.106 | 0.204 | 0.346 | 0.033 | 0.223 | 0 | - | - | 0.0832 | - | - | - |

| | | | | | | | | | | | | | |
|---|---|---|---|---|---|---|---|---|---|---|---|---|---|
| 38 | 0.249 | 0.194 | 0.017 | - | - | 0.206 | 0 | - | - | 0.0868 | 0.0225 | 1.05E-15 | - |
| 26 | 0.293 | 0.113 | 0.204 | 0.034 | - | 0.220 | 0 | - | - | 0.0809 | - | - | - |
| 24 | 0.252 | 0.197 | 0.038 | - | - | 0.204 | 0 | - | - | 0.0694 | - | - | - |
| 32 | 0.311 | 0.075 | 0.208 | 0.020 | - | 0.211 | 0 | 0.288 | - | 0.0911 | - | - | - |
| 44 | 0.267 | 0.112 | 0.021 | - | - | 0.228 | 0 | 0.176 | - | 0.0867 | 8.22E-15 | - | - |
| 30 | 0.245 | 0.084 | - | - | - | 0.224 | 0 | - | - | 4.88E-09 | 0.1281 | 0.0547 | 0.0285 |
| 42 | 0.176 | 0.111 | 0.250 | - | - | 0.139 | 0 | 0.238 | - | 0.0941 | 0.0176 | - | - |
| 36 | 0.306 | 0.081 | 0.145 | - | - | 0.227 | 0 | - | - | 0.1030 | 0.0268 | 1.50E-15 | - |
| 34 | 0.302 | 0.096 | 0.149 | - | - | 0.205 | 0 | 0.356 | 0.164 | 0.0804 | - | - | - |
| 4 | 0.212 | - | - | - | - | 0.205 | 0 | - | - | 0.0794 | 1.37E-15 | - | - |
| 6 | 0.233 | 0.118 | - | - | - | 0.217 | 0 | - | - | 0.0876 | 5.00E-15 | - | - |
| 20 | 0.212 | - | - | - | - | 0.205 | 0 | - | - | 0.0883 | 0.0251 | 0.0266 | - |
| 22 | 0.232 | 0.122 | - | - | - | 0.218 | 0 | - | - | 0.0982 | 0.0268 | 0.0284 | - |
| 12 | 0.212 | - | - | - | - | 0.185 | 0 | 0.300 | - | 0.0693 | - | - | - |
| 14 | 0.212 | - | - | - | - | 0.204 | 0 | - | - | 0.0266 | - | - | - |
| 1 | 0.213 | - | - | - | - | 0.205 | 0 | - | - | 0.0696 | - | - | - |
| 18 | 0.234 | 0.115 | - | - | - | 0.122 | 0 | 0.248 | 0.242 | 0.0746 | - | - | - |
| 2 | 0.227 | 0.140 | - | - | - | 0.213 | 0 | - | - | 0.0748 | - | - | - |
| 8 | 0.250 | 0.101 | 0.206 | - | - | 0.201 | 0 | 0.320 | - | 0.0848 | 0.0369 | - | - |
| 16 | 0.196 | 0.265 | 0.111 | - | - | 0.219 | 0 | - | - | 0.0786 | - | - | - |
| 10 | 0.216 | 0.198 | - | - | - | 0.204 | 0 | - | - | 0.0695 | - | - | - |
| 40 | 0.195 | 0.112 | 0.265 | - | - | 0.208 | 0 | 0.226 | - | 0.0782 | - | - | - |

**Table S4**. Model comparison based on AIC for the models tested with the extant-species tree. Models differ in what state changes are allowed, and whether state changes occur during speciation. CR = Constant Rates; CTD = Concealed trait-dependent; ETD = Examined trait-dependent; K = number of free parameters.

| Model | Trait-dependence | loglikelihood | K | AIC | ΔAIC | AICw |
|---|---|---|---|---|---|---|
| 30 | ETD | -144.302 | 7 | 304.605 | 0 | 0.365 |
| 22 | ETD | -145.778 | 6 | 305.556 | 0.951 | 0.227 |
| 38 | ETD | -145.035 | 7 | 306.070 | 1.466 | 0.176 |
| 21 | CTD | -147.864 | 5 | 307.727 | 3.123 | 0.077 |
| 36 | ETD | -146.216 | 7 | 308.431 | 3.827 | 0.054 |
| 23 | CTD | -147.746 | 6 | 309.492 | 4.888 | 0.032 |
| 20 | ETD | -149.287 | 5 | 310.574 | 5.969 | 0.018 |
| 31 | CTD | -147.707 | 7 | 311.414 | 6.809 | 0.012 |
| 26 | ETD | -149.490 | 6 | 312.979 | 8.375 | 0.006 |
| 37 | CTD | -148.750 | 7 | 313.500 | 8.896 | 0.004 |
| 2 | ETD | -151.964 | 4 | 313.928 | 9.323 | 0.003 |
| 39 | CTD | -148.999 | 7 | 313.998 | 9.393 | 0.003 |
| 6 | ETD | -151.149 | 5 | 314.297 | 9.693 | 0.003 |
| 16 | ETD | -151.175 | 5 | 314.351 | 9.746 | 0.003 |
| 44 | ETD | -149.223 | 7 | 314.446 | 9.841 | 0.003 |
| 28 | ETD | -149.460 | 7 | 314.920 | 10.315 | 0.002 |
| 32 | ETD | -149.490 | 7 | 314.979 | 10.375 | 0.002 |
| 24 | ETD | -151.539 | 7 | 315.078 | 10.473 | 0.002 |
| 13 | CTD | -152.976 | 4 | 315.952 | 11.347 | 0.001 |
| 40 | ETD | -151.175 | 6 | 316.351 | 11.746 | 0.001 |
| 18 | ETD | -151.605 | 6 | 317.209 | 12.605 | 0.001 |
| 5 | CTD | -153.638 | 4 | 317.277 | 12.672 | 0.001 |
| 42 | ETD | -150.897 | 7 | 317.794 | 13.189 | 0.0 |
| 8 | ETD | -150.903 | 7 | 317.805 | 13.201 | 0.0 |
| 19 | CTD | -151.955 | 6 | 317.909 | 13.305 | 0.0 |
| 11 | CTD | -154.347 | 4 | 318.694 | 14.090 | 0.0 |
| 3 | CTD | -154.366 | 4 | 318.733 | 14.128 | 0.0 |
| 15 | CTD | -154.410 | 4 | 318.820 | 14.216 | 0.0 |
| 34 | ETD | -151.466 | 7 | 318.932 | 14.328 | 0.0 |
| 1 | CR | -155.517 | 3 | 319.035 | 14.430 | 0.0 |
| 7 | CTD | -153.701 | 5 | 319.403 | 14.798 | 0.0 |
| 4 | ETD | -154.977 | 4 | 319.953 | 15.349 | 0.0 |
| 33 | CTD | -152.023 | 7 | 320.046 | 15.441 | 0.0 |
| 43 | CTD | -152.108 | 7 | 320.215 | 15.611 | 0.0 |
| 9 | CTD | -152.161 | 7 | 320.322 | 15.717 | 0.0 |
| 14 | ETD | -155.244 | 4 | 320.488 | 15.883 | 0.0 |
| 35 | CTD | -152.292 | 7 | 320.583 | 15.979 | 0.0 |
| 25 | CTD | -154.317 | 5 | 320.634 | 16.029 | 0.0 |
| 17 | CTD | -154.348 | 5 | 320.697 | 16.092 | 0.0 |
| 12 | ETD | -155.445 | 4 | 320.891 | 16.286 | 0.0 |
| 41 | CTD | -153.512 | 6 | 321.024 | 16.419 | 0.0 |
| 10 | ETD | -155.515 | 4 | 321.029 | 16.425 | 0.0 |
| 45 | CTD | -152.766 | 7 | 321.532 | 16.928 | 0.0 |
| 27 | CTD | -154.341 | 6 | 322.682 | 18.077 | 0.0 |
| 29 | CTD | -154.229 | 7 | 324.458 | 19.854 | 0.0 |

**Table S5.** Estimates of all the parameters used in each model for the extant-species tree. Models are ordered from best fitted model to the worst.

| Model | $\lambda_1$ | $\lambda_2$ | $\lambda_3$ | $\lambda_4$ | $\lambda_5$ | $\mu_1$ | $\mu_2$ | $\mu_3$ | $\mu_4$ | $q_1$ | $q_2$ | $q_3$ | $q_4$ |
|---|---|---|---|---|---|---|---|---|---|---|---|---|---|
| 30 | 0.109 | 0.361 | - | - | - | 1.698E-08 | 0 | - | - | 0.0384 | 0.126 | 0.031 | 5.14E-16 |
| 22 | 0.101 | 0.384 | - | - | - | 1.882E-09 | 0 | - | - | 0.1004 | 0.029 | 1.00E-16 | - |
| 38 | 0.190 | 0.136 | 0.021 | - | - | 8.459E-08 | 0 | - | - | 0.1016 | 4.605E-08 | 2.60E-17 | - |
| 21 | 0.076 | - | - | - | - | 0.137 | 0 | - | - | 0.1424 | 0.038 | 3.72E-16 | - |
| 36 | 0.137 | 0.306 | 0.091 | - | - | 3.122E-08 | 0 | - | - | 0.1019 | 5.924E-16 | 2.04E-15 | - |
| 23 | 0.078 | 0.194 | - | - | - | 0.150 | 0 | - | - | 0.1438 | 0.039 | 2.28E-16 | - |
| 20 | 0.173 | - | - | - | - | 0.069 | 0 | - | - | 0.1172 | 0.037 | 5.19E-16 | - |
| 31 | 0.079 | 0.190 | - | - | - | 0.122 | 0 | - | - | 0.1016 | 0.154 | 0.039 | 6.65E-17 |
| 26 | 0.131 | 0.364 | 0.136 | 0.025 | - | 1.649E-15 | 0 | - | - | 0.0635 | - | - | - |
| 37 | 0.077 | 0.192 | 0.184 | - | - | 0.130 | 0 | - | - | 0.1148 | 1.370E-09 | 0.146 | - |
| 2 | 0.105 | 0.370 | - | - | - | 9.011E-16 | 0 | - | - | 0.0648 | - | - | - |
| 39 | 0.078 | 0.198 | 0.186 | - | - | 0.119 | 0 | - | - | 0.1146 | 1.232E-08 | 0.133 | - |
| 6 | 0.106 | 0.374 | - | - | - | 7.860E-10 | 0 | - | - | 0.0700 | 0.003 | - | - |
| 16 | 0.132 | 0.071 | 0.383 | - | - | 9.125E-09 | 0 | - | - | 0.0672 | - | - | - |
| 44 | 0.137 | 0.290 | 0.021 | - | - | 6.138E-15 | 0 | 5.839E-08 | - | 0.0716 | 7.172E-18 | - | - |
| 28 | 0.118 | 0.362 | 0.136 | 0.140 | 0.024 | 5.967E-09 | 0 | - | - | 0.0633 | - | - | - |
| 32 | 0.131 | 0.364 | 0.136 | 0.025 | - | 5.691E-14 | 0 | 1.265E-08 | - | 0.0635 | - | - | - |
| 24 | 0.197 | 0.137 | 0.024 | - | - | 1.327E-07 | 0 | - | - | 0.0627 | 0.031 | 1.83E-16 | - |
| 13 | 0.077 | - | - | - | - | 0.243 | 0 | 1.130E-09 | - | 0.1004 | - | - | - |
| 40 | 0.132 | 0.383 | 0.071 | - | - | 7.638E-10 | 0 | 2.615E-14 | - | 0.0672 | - | - | - |
| 18 | 0.110 | 0.384 | - | - | - | 5.320E-07 | 0 | 2.450E-09 | 0.035 | 0.0666 | - | - | - |
| 5 | 0.076 | - | - | - | - | 0.134 | 0 | - | - | 0.0911 | 3.816E-10 | - | - |
| 42 | 0.114 | 0.378 | 0.103 | - | - | 6.034E-08 | 0 | 1.442E-07 | - | 0.0749 | 0.024 | - | - |
| 8 | 0.081 | 0.389 | 0.136 | - | - | 1.155E-15 | 0 | 0.066 | - | 0.0776 | 0.028 | - | - |

| | | | | | | | | | | | | | |
|---|---|---|---|---|---|---|---|---|---|---|---|---|---|
| 19 | 0.074 | 0.173 | - | - | - | 0.548 | 0 | 0.000 | 0.405 | 0.1004 | - | - | - |
| 11 | 0.081 | 0.190 | - | - | - | 0.122 | 0 | - | - | 0.0867 | - | - | - |
| 3 | 0.079 | 0.190 | - | - | - | 0.118 | 0 | - | - | 0.0866 | - | - | - |
| 15 | 0.082 | - | - | - | - | 0.106 | 0 | - | - | 0.0885 | 0.045 | - | - |
| 34 | 0.128 | 0.569 | 0.085 | - | - | 8.315E-07 | 0 | 0.235 | 4.462E-09 | 0.0841 | - | - | - |
| 1 | 0.166 | - | - | - | - | 0.053 | 0 | - | - | 0.0728 | - | - | - |
| 7 | 0.078 | 0.177 | - | - | - | 0.124 | 0 | - | - | 0.0899 | 1.284E-15 | - | - |
| 4 | 0.166 | - | - | - | - | 0.051 | 0 | - | - | 0.0765 | 6.752E-15 | - | - |
| 33 | 0.079 | 0.169 | 0.168 | 0.167 | - | 0.427 | 0 | 4.370E-09 | - | 0.0970 | - | - | - |
| 43 | 0.078 | 0.188 | 0.182 | - | - | 0.716 | 0 | 0.112 | - | 0.1328 | 0.034 | - | - |
| 9 | 0.078 | 0.175 | 0.174 | - | - | 0.261 | 0 | 1.408E-07 | - | 0.1125 | 0.041 | - | - |
| 14 | 0.166 | - | - | - | - | 0.052 | 0 | - | - | 0.0765 | 0.038 | - | - |
| 35 | 0.071 | 0.179 | 0.173 | - | - | 0.329 | 0 | 7.367E-11 | 0.306 | 0.1060 | - | - | - |
| 25 | 0.076 | 0.191 | 0.181 | - | - | 0.123 | 0 | - | - | 0.0864 | - | - | - |
| 17 | 0.076 | 0.190 | 0.182 | - | - | 0.118 | 0 | - | - | 0.0862 | - | - | - |
| 12 | 0.163 | - | - | - | - | 0.052 | 0 | 1.599E-14 | - | 0.0707 | - | - | - |
| 41 | 0.073 | 0.177 | 0.173 | - | - | 0.099 | 0 | 0.204 | - | 0.0886 | - | - | - |
| 10 | 0.167 | 0.163 | - | - | - | 0.052 | 0 | - | - | 0.0727 | - | - | - |
| 45 | 0.072 | 0.175 | 0.170 | - | - | 0.256 | 0 | 9.252E-08 | - | 0.0946 | 2.225E-16 | - | - |
| 27 | 0.079 | 0.189 | 0.180 | 0.178 | - | 0.117 | 0 | - | - | 0.0860 | - | - | - |
| 29 | 0.076 | 0.187 | 0.187 | 0.175 | 0.176 | 0.121 | 0 | - | - | 0.0869 | - | - | - |

### FIGURE LIST

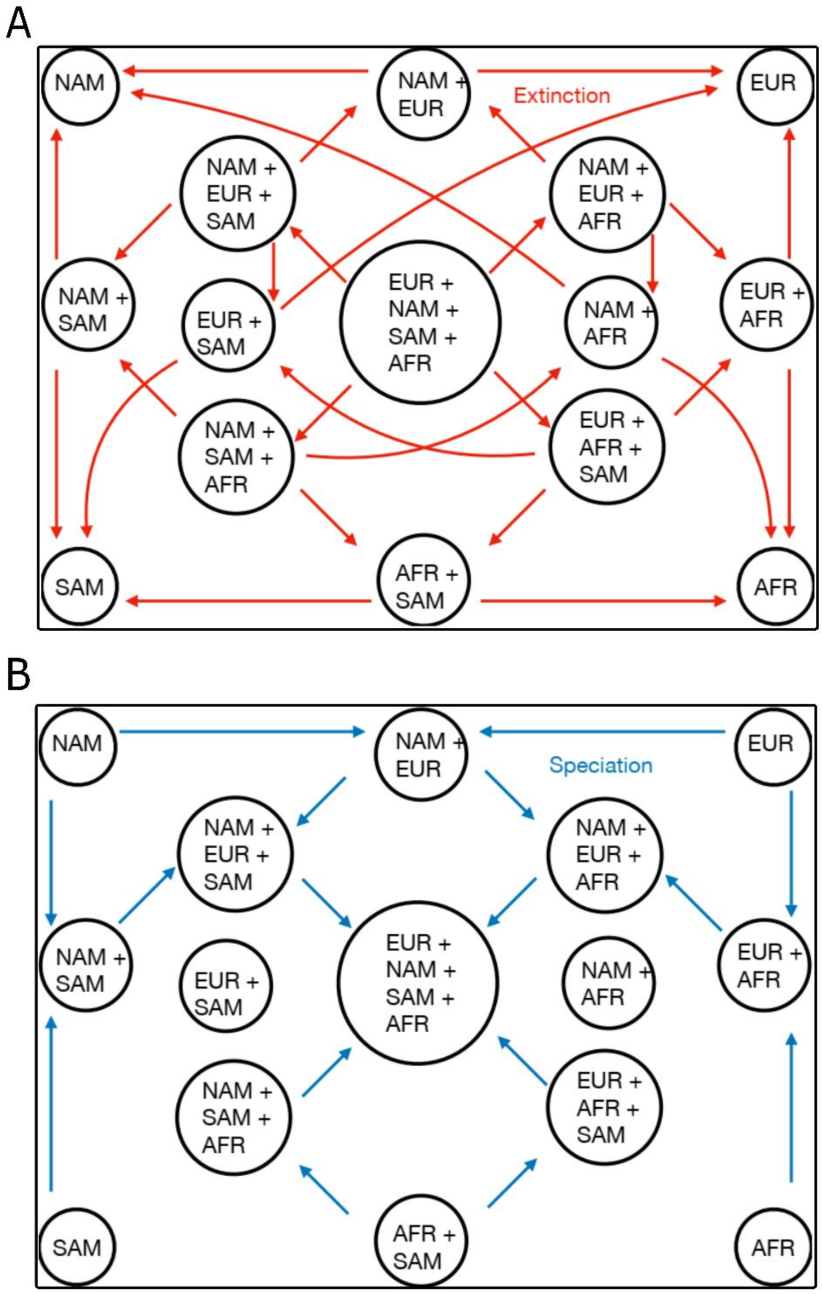


**Figure S1.** Graphical representation of all possible ways species can go extinct (A) or speciate (B) in our 45 models.
